# When the whole is only the parts: non-holistic object parts predominate face-cell responses to illusory faces

**DOI:** 10.1101/2023.09.22.558887

**Authors:** Saloni Sharma, Kasper Vinken, Margaret S. Livingstone

**Author notes:** **Author contributions**: Conceptualization: S.S., K.V., and M.S.L. Investigation: S.S. Formal analysis: S.S and K.V. Methodology: S.S and K.V. Visualization: S.S and K.V. Supervision: K.V. and M.S.L. Writing—original draft: S.S. Writing—review and editing: S.S., K.V., and M.S.L. **Competing interests**: The authors declare that they have no competing interests.

## Abstract

Humans are inclined to perceive faces in everyday objects with a face-like configuration. This illusion, known as face pareidolia, is often attributed to a specialized network of ‘face cells’ in primates. We found that face cells in macaque inferotemporal cortex responded selectively to pareidolia images, but this selectivity did not require a holistic, face-like configuration, nor did it encode human faceness ratings. Instead, it was driven mostly by isolated object parts that are perceived as eyes only within a face-like context. These object parts lack usual characteristics of primate eyes, pointing to the role of lower-level features. Our results suggest that face-cell responses are dominated by local, generic features, unlike primate visual perception, which requires holistic information. These findings caution against interpreting neural activity through the lens of human perception. Doing so could impose human perceptual biases, like seeing faces where none exist, onto our understanding of neural activity.

## Introduction

The primary goal of vision neuroscience is to understand the link between neural activity in the brain and visual perception. A central approach is to interpret what each neuron “likes to see”, to link the role of individual neurons back to perception. The groundwork for this notion of interpretable neurons was laid by Hubel and Wiesel (Hubel & Wiesel, 1962, 1974), who suggested that cells at the early stages of the visual system encode basic features like edges and orientations. At higher levels of visual processing, the discovery of neurons with selective responses for complex visual stimuli such as faces, hands, or objects in the inferotemporal (IT) cortex(Desimone et al., 1984; Desimone & Gross, 1979; Gross et al., 1972; Perrett et al., 1982; K. Tanaka et al., 1991) led to the suggestion that these neurons directly encode the perception of high-level categories (Desimone & Gross, 1979; Taubert et al., 2020). A challenge with the approach of interpreting what a neuron responds to, is that it is inherently subject to the biases of human perception itself. That is, when a neuron responds to faces, it may not respond to what we as humans see in those images.

Decades of psychophysical experiments suggest that humans and other primates perceive faces holistically (Maurer et al., 2002; Parr, 2011; J. W. Tanaka & Simonyi, 2016). For instance, humans recognize face identities more easily when the parts are embedded in the context of a whole face (whole-parts effect; J. W. Tanaka & Farah, 1993) and when faces are presented upright rather than inverted (Yin, 1969). The dependence of face perception on holistic context was also demonstrated effectively with the Thatcher illusion (Thompson, 1980), where humans and macaques recognize changes in the orientation of facial features (such as eyes) only in the context of an upright face, and not in an inverted(Dahl et al., 2010). At the neural level, extensive research has linked face perception to the activity of face cells in IT cortex. Face cells respond more to faces than to non-face objects (Bruce et al., 1981; Gross et al., 1972; Tsao et al., 2006) and have been causally implicated in the perception of faces (A. Afraz et al., 2015; S.-R. Afraz et al., 2006; Azadi et al., 2023; Moeller et al., 2017; Sadagopan et al., 2017). Several studies have suggested hallmark properties of configural or holistic face processing in the responses of individual face cells. For instance, scrambling faces into many smaller parts significantly reduces the firing rate, which suggests a holistic dependence on facial configuration (Desimone et al., 1984; Perrett et al., 1982). Further, face cells responses are reduced to inverted faces (face inversion effect; Freiwald et al., 2009; Taubert et al., 2015b) or when the eyes in an upright face are inverted (thatcher illusion; Taubert et al., 2015a). On the other hand, it is less clear if face cells demonstrate the whole-parts effect. One study showed that the gain of tuning in face cells for a particular feature was twice as high when presented in combination with other features in a face-like arrangement (Freiwald et al., 2009; Hesse & Tsao, 2020). However, other studies showed that face cells respond to isolated face parts without requiring the context of a face(Issa & DiCarlo, 2012; Waidmann et al., 2022). One potential interpretation for these conflicting findings is that the studies with responses to isolated features used real face parts, which are recognizable as such and may imply the presence of an occluded face(Arcaro, Ponce, et al., 2020). Yet, there is also evidence showing that face cells reliably respond to nonface objects, which usually lack a face-like configuration or recognizable face parts (Meyers et al., 2015; Vinken et al., 2023). Thus, while holistic processing plays a role in face perception, its role in the activity of face cells remains less clear. The question is further complicated by studies that used real face parts, which confound the interpretation of whether and when face cells require holistic information.

Interestingly, there are examples where humans do not need actual faces or facial features to perceive a face, such as “Jesus in toast” or “Man in the moon”. The tendency to see illusory faces in inanimate objects is referred to as face pareidolia. Various studies have shown that such illusory faces are perceived as highly face-like (Hadjikhani et al., 2009; Omer et al., 2019; Taubert et al., 2017; Wardle et al., 2020, 2022) and, importantly, that they engage face selective regions in both humans and monkeys (Akdeniz et al., 2018; Decramer et al., 2021; Liu et al., 2014; Taubert et al., 2020, 2022; Wardle et al., 2020). The perception of illusory faces has been attributed to the configural processing of individual object parts that are spuriously arranged in a face-like configuration (Hadjikhani et al., 2009; Omer et al., 2019; Taubert et al., 2017; Wardle et al., 2020, 2022). The importance of the configuration is stressed by the fact that normal facial features are absent in these images and are replaced by notably heterogeneous features clearly incongruent with real faces, such as the rocky surface of the moon (Taubert et al., 2017). Indeed, an affinity for some aspects of a face-like configuration is present early in development and suggested to facilitate rapid detection, even when normal facial features are absent (Caruana & Seymour, 2021; Fantz, 1963; Goren et al., 1975; Johnson et al., 1991; Keys et al., 2021; Kuwahata et al., 2004; Maurer et al., 2002; Mondloch et al., 1999; Parr, 2011; Reid et al., 2017; Tsao & Livingstone, 2008; Valenza et al., 1996). Taken together, it seems parsimonious to interpret neural selectivity for pareidolic images, defined by a face-like configuration, as evidence for holistic neural processing that mirrors perception. However, for such images that are inherently subject to a perceptual bias, it may be particularly challenging to interpret what a neuron within the face-selective regions “likes to see” in them.

Here, we address the question whether the face-like configuration is critical in driving face cell responses to illusory faces. We recorded multiunit activity in central (CIT) and anterior IT (AIT) in 8 monkeys (n = 4 for CIT, n = 5 for AIT). Specifically, we investigated 1) whether macaque face cells selectively respond to objects with an illusory face compared to those without, 2) if a face-like configuration is necessary for driving pareidolia selectivity, 3) what other, non-holistic attributes drive pareidolia selectivity, and 4) if face-cell responses reflect how face-like an object looks to a human observer. Overall, we found that face-selective units in IT, which responded more to faces than to non-faces, were also pareidolia selective, in that they responded more to pareidolia images than to matched controls. We found a strong positive correlation between face selectivity and pareidolia selectivity in both central and anterior IT. Strikingly, eliminating the face-like configuration by quadrant scrambling the pareidolia images did not abolish pareidolia selectivity of face cells, or the correlation between face and pareidolia selectivity. When we presented the four quadrants in isolation, we found that the pareidolia selectivity of face-selective units was driven primarily by the quadrants containing the object features that act as eyes of the illusory face, which we will refer to as pareidolia eye features throughout the paper, for brevity. Further, we found that pareidolia selectivity was predicted by neural tuning estimated from non-face, non-pareidolia images. Finally, we found that face-cell activity to pareidolia images could not be explained by human faceness ratings, and faceness ratings could only weakly be predicted from a linear combination of the neural responses. Taken together, these results suggest that face cells do not respond to pareidolia images for the same reasons we see faces in them. They responded mostly to non-holistic object features rather than a face-like configuration or object faceness per se. Thus, we show that individual face-cell responses to pareidolic images do not mirror the holistic perceptual experience of an illusory face.

## Results

In our main experiment, we presented 100 pareidolia images and 100 control images to 8 macaque monkeys in a Rapid Serial Visual Presentation (rsvp)-style paradigm. The pareidolia images were images of objects that evoked the illusion of a face, whereas control images were matched in object identity to the pareidolia images but did not evoke an illusory face experience (Taubert et al., 2017; Wardle et al., 2022). We recorded neural activity from central (CIT) and anterior IT (AIT) while these images were presented. Additionally, we presented quadrant scrambled pareidolia and control images, in which each image was divided into four quadrants and the quadrants were shuffled. This ensured that the individual features were preserved but the images did not retain the global T-shaped configuration and thus did not evoke the perceptual experience of a face. In a second experiment, we presented only scrambled versions of pareidolia images that the monkeys had never seen before. In a third experiment, we presented the four quadrants of the original and scrambled configurations in isolation. Finally, in our final experiment we presented the monkeys with 200 pareidolia images that had been given faceness ratings on an 11-point scale (0 “cannot see a face” to 10 “easily see a face”) by human subjects in a recent study (Wardle et al., 2022).

### Units in CIT and AIT show pareidolia selectivity

We recorded multiunit spiking activity in CIT (n=208 sites, pooled across 4 macaques) and AIT (n=163 sites, pooled across 5 macaques). Two of the macaques included in this study were deprived of seeing faces for the first year of their life. However, at the time these experiments were run they both had more than two years’ experience with faces. We used 40 faces, 40 non-faces, 100 pareidolia images and 100 matched control objects (see examples in Fig. 1a). We found that, on average, the recorded populations contained a range of face selective (responded more to faces than nonfaces) and pareidolia selective (responded more to pareidolia images than to matched controls) units in CIT (Fig 1. b, c) and AIT (Fig. 1g, f). In Fig. 1d, h, we show some example units which showed some relation between face selectivity and pareidolia selectivity. Next, we investigated whether the magnitude of a unit’s face selectivity also predicted the magnitude of its pareidolia selectivity.

**Fig. 1.**
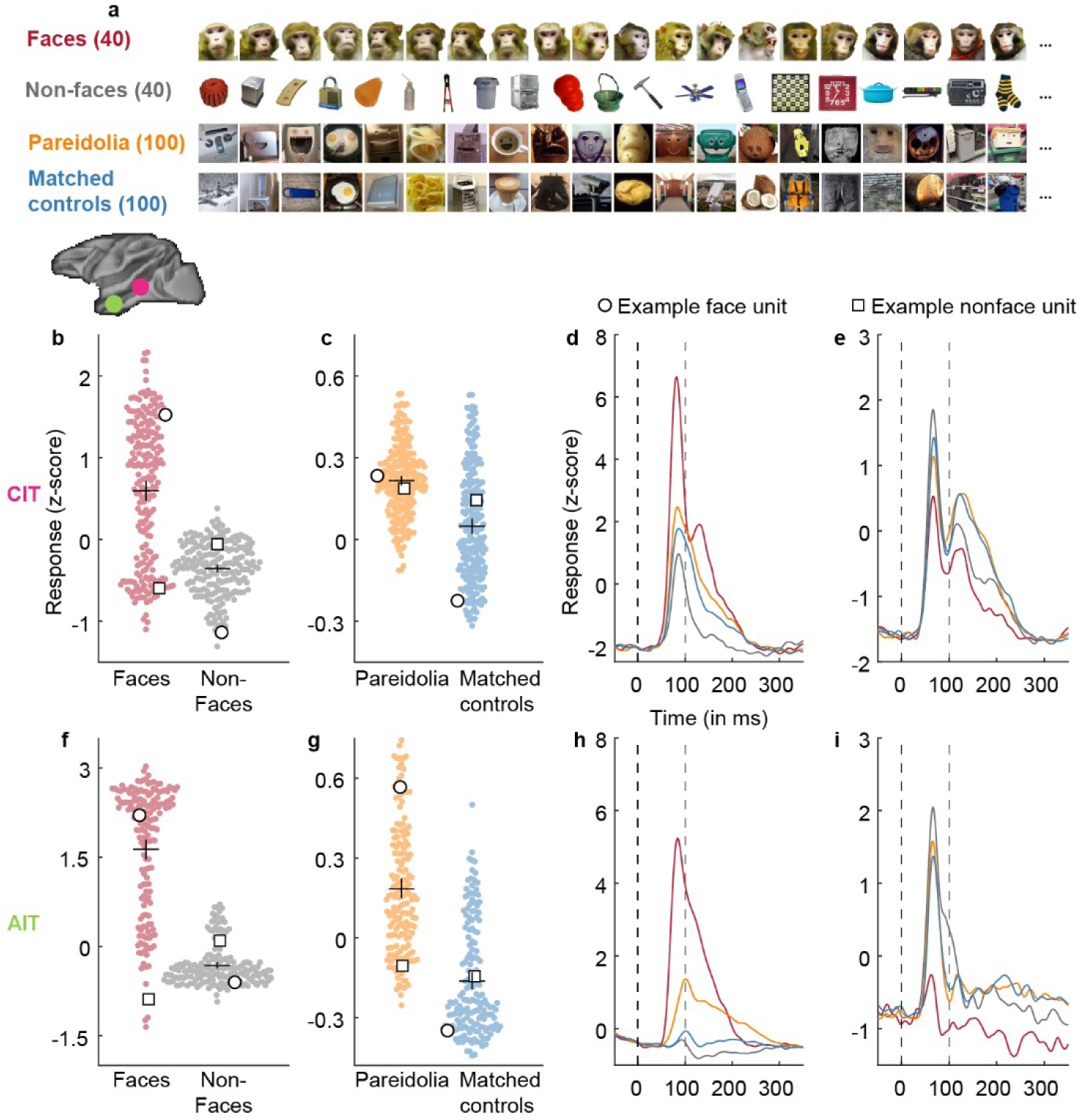
Units in CIT and AIT show pareidolia selectivity. **a.** Example visual stimuli used in the experiment. Faces and non-face objects were included to be able to calculate the face selectivity of the neural units. Each pareidolia image included a matched control object, which has the same object identity, but does not invoke the perceptual experience of a face. **b.** The beeswarm plot shows normalized neural response per unit averaged across 150ms time window starting at response onset (see Methods) to faces (red) and non-faces (grey) in central IT (CIT). Open circles and squares represent the example face (circle) and nonface (square) units shown in d, e. **c**. Plot showing normalized neural response per unit to pareidolia (orange) and matched control objects (blue). Same conventions as in b. **d**. The time course of an example face unit (depicted by an open circle in b, c) in CIT. X-axis represents time in milliseconds and y-axis shows the normalized neural response. Vertical dashed line at t = 0 indicates stimulus onset. Grey line at t = 100 indicates stimulus offset. **e.** Time course of an example nonface unit (depicted by an open square in b, c). Same conventions as in d. **f, g.** Beeswarm plots showing normalized neural responses per unit anterior IT (AIT). Same conventions as in b, c. **h, i.** Time course of an example face unit (h) and nonface unit (i) in AIT. Same conventions as in d, e. The small inset of the macaque brain shows the approximate location of the recording sites in CIT (pink circle) and AIT (green circle).

### Pareidolia selectivity is correlated with face selectivity

We quantified category selectivity by computing a d’ selectivity index for faces and for pareidolias for each unit. This index reflects the difference in mean responses between two image categories in standard deviation units (for instance, a face d’ > 0 indicates units respond more to faces than non-faces; pareidolia d’ > 0 indicates units respond more to pareidolia than to matched controls), which we report in all further analysis. Further, we use the term “face units” to refer to units with a face d’>1 and “nonface units” to refer to those with a face d’<1. The average pareidolia d’ for face units was 0.3 ± 0.16 (t_112_ = 19.36, p = 1.6 x 10^-37^) whereas for non-face units it was 0.04 ± 0.16 (t_94_ = 2.3, p = 0.03) in CIT. In AIT, the average pareidolia d’ for face units was 0.53 ± 0.32 (t_121_ = 17.98, p = 5.52 x 10^-36^), whereas for non-face units it was 0.3 ± 0.2 (t_40_ = 9.2, p = 2.01 x 10^-11^). Note that these averages are lower than the threshold face d’ value of 1 used to select face-selective units, which is expected given the overall lower response to pareidolia images (see Fig. 1). In both regions, the average pareidolia d’ was significantly higher for face units than for non-face units (CIT: t_206_ = 11.36, p = 1.56 x 10^-23^, AIT: t_161_ = 4.38, p = 2.15 x 10^-5^). The pareidolia and face selectivity of each unit are shown in Fig. 2a, b, which shows that pareidolia selectivity is positively correlated with face selectivity (Fig. 2a, b; CIT: n = 208, Pearson’s r = 0.7, p = 9.39 x 10^-32^, AIT: n = 163, Pearson’s r = 0.61, p = 7.62 x 10^-18^). This positive correlation between face and pareidolia selectivity was true for each of the 8 monkeys individually (CIT, n = 4, AIT, n = 5; Table S1). One-sample t-tests on the Fischer-transformed correlation values indicate that this correlation was significant across monkeys (CIT: t_3_ = 4.62, p = 0.0096; AIT: t_4_ = 8.19, p = 0.00061). This means that a unit’s face selectivity and its pareidolia selectivity are closely related in IT cortex and could both be driven by image attributes shared between faces and pareidolia images. It is worth noting that, although we divided units into face and non-face units, both face selectivity and pareidolia selectivity followed a graded continuum, rather than a discrete step function.

**Fig. 2.**
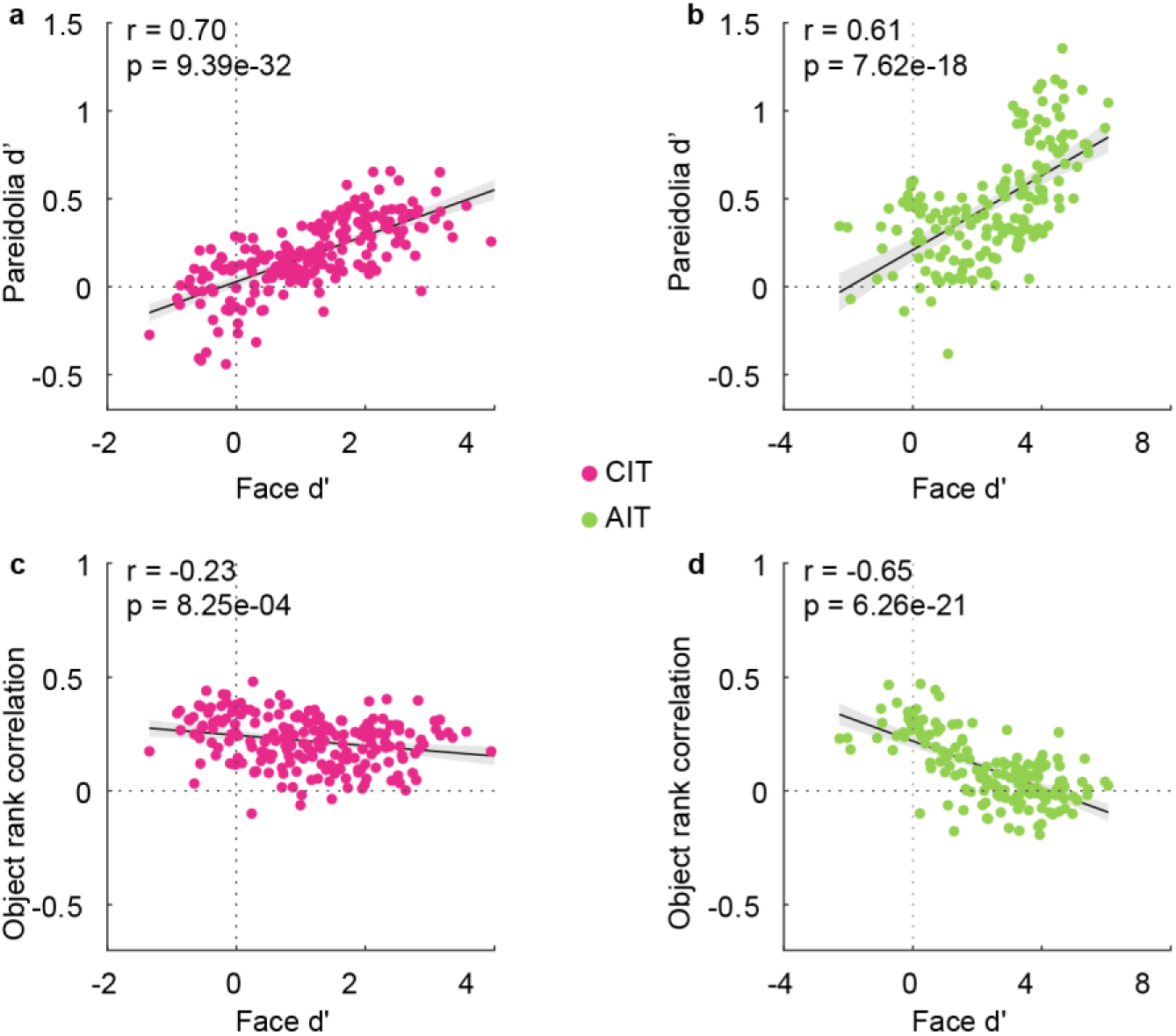
Pareidolia selectivity is correlated with face selectivity. **a.** Scatterplot showing the correlation between face selectivity (face d’) plotted on the x-axis and pareidolia selectivity (pareidolia d’) on the y-axis. Each dot depicts a neural unit in CIT (n = 208, pooled across 4 monkeys). The black line indicates an ordinary least squares (OLS) linear regression fit, with shaded 95% confidence bounds. **b.** Scatterplot showing the correlation between face and pareidolia selectivity in AIT (n = 163, pooled across 5 monkeys). Same conventions as in (a). **c, d.** Scatterplot showing the correlation of face selectivity to object rank correlation in CIT (c) and AIT (d).

For each pareidolia image, we had one matched control image of the same kind of object (e.g., a chair or a faucet). To evaluate whether the selectivity profile across various objects was the same for pareidolia and control images, we calculated an object rank correlation for each unit. This metric was defined as the Spearman correlation between the response vector to all pareidolia images and the response vector to all corresponding controls. A positive object rank correlation indicates that a neural unit responds to features shared between pareidolia images and their matched controls, but distinct across object identities (e.g., properties like texture or color that are associated with object identity in our stimulus set). Conversely, an object rank correlation of zero indicates that the unit did not selectively respond to features differentiating between object identities. We found a significant positive object rank correlation in face units (CIT: 0.2 ± 0.1, t_112_ = 21.43, p = 2 x 10^-41^, AIT = 0.04 ± 0.1, t_121_ = 3.86, p = 1.82 x 10^-4^) and in non-face units (CIT: 0.25 ± 0.12, t_94_ = 20.5, p = 1.93 x 10^-36^, AIT = 0.27 ± 0.1, t_40_ = 15.67, p = 1.14 x 10^-18^). Further, the object rank correlation for face units was significantly lower than for nonface units, particularly in AIT (CIT: t_206_ = −3.23, p = 0.0014, AIT: t_161_ = −12.44, p = 2.46 x 10^-25^). This effect is explained by the negative correlation between face selectivity and object rank correlation (Fig. 2c, d; CIT: n = 208, Pearson’s r = - 0.23, p = 0.00083, AIT: n = 163, Pearson’s r = −0.65, p = 6.26 x 10^-21^). Taken together, this result indicates that the more face selective units in CIT and AIT were driven less by object features that differentiate between object identities, and more by object features shared between actual and illusory faces. This raises the question: what specific image properties caused face cells to respond selectively to the pareidolia images? Are these the same properties that instigate the illusory perception of faces in these images, that is, the face-like configuration?

### Quadrant scrambling preserves the relationship between pareidolia and face selectivity

A critical factor that seems to drive the illusory perception of a face in a pareidolia image is the global configuration or the spatial arrangement of certain object parts, such as circular spots or curved edges. When arranged in the T-shaped arrangement of features typically associated with faces, these object parts appear as two eyes above a mouth, thus evoking the perceptual experience of an illusory face (see examples in Fig. 3b, top). This face-like global configuration has been suggested to facilitate rapid face detection through template matching, even in the absence of normal facial features (Caruana & Seymour, 2021; Keys et al., 2021; Maurer et al., 2002; Palmer & Clifford, 2020; Tsao & Livingstone, 2008; Zhou et al., 2021). Further, the preference for the face-like configuration over a scrambled configuration, even when composed of simple shapes such as circles or squares (such as in our cartoon example in Fig.3a), is already present in early infancy and continues into adulthood for both humans and monkeys (Fantz, 1963; Goren et al., 1975; Johnson et al., 1991; Keating & Keating, 1982; Kuwahata et al., 2004; Mondloch et al., 1999; Reid et al., 2017; Tomalski et al., 2009; Valenza et al., 1996). Moreover, the face-like configuration, but not the scrambled configuration, has been shown to interfere with local feature detection or peripheral face recognition, suggesting that we find such a configuration difficult to ignore (Sun & Balas, 2015; Suzuki & Cavanagh, 1995). Thus, collectively, extensive prior research underscores the significance of the global face-like configuration in face perception. Indeed, this significance also becomes apparent in the cartoon example in Fig. 3a: the perception of a face is strong when two dots are horizontally aligned and above a horizontal line. Yet, the illusion of a face is eliminated when the face-like configuration is removed by scrambling the configuration (See cartoon example in Fig. 3a and scrambled pareidolia examples in Fig. 3b, bottom row). Would removing the face-like configuration similarly eliminate the pareidolia selectivity of face cells?

**Fig. 3.**
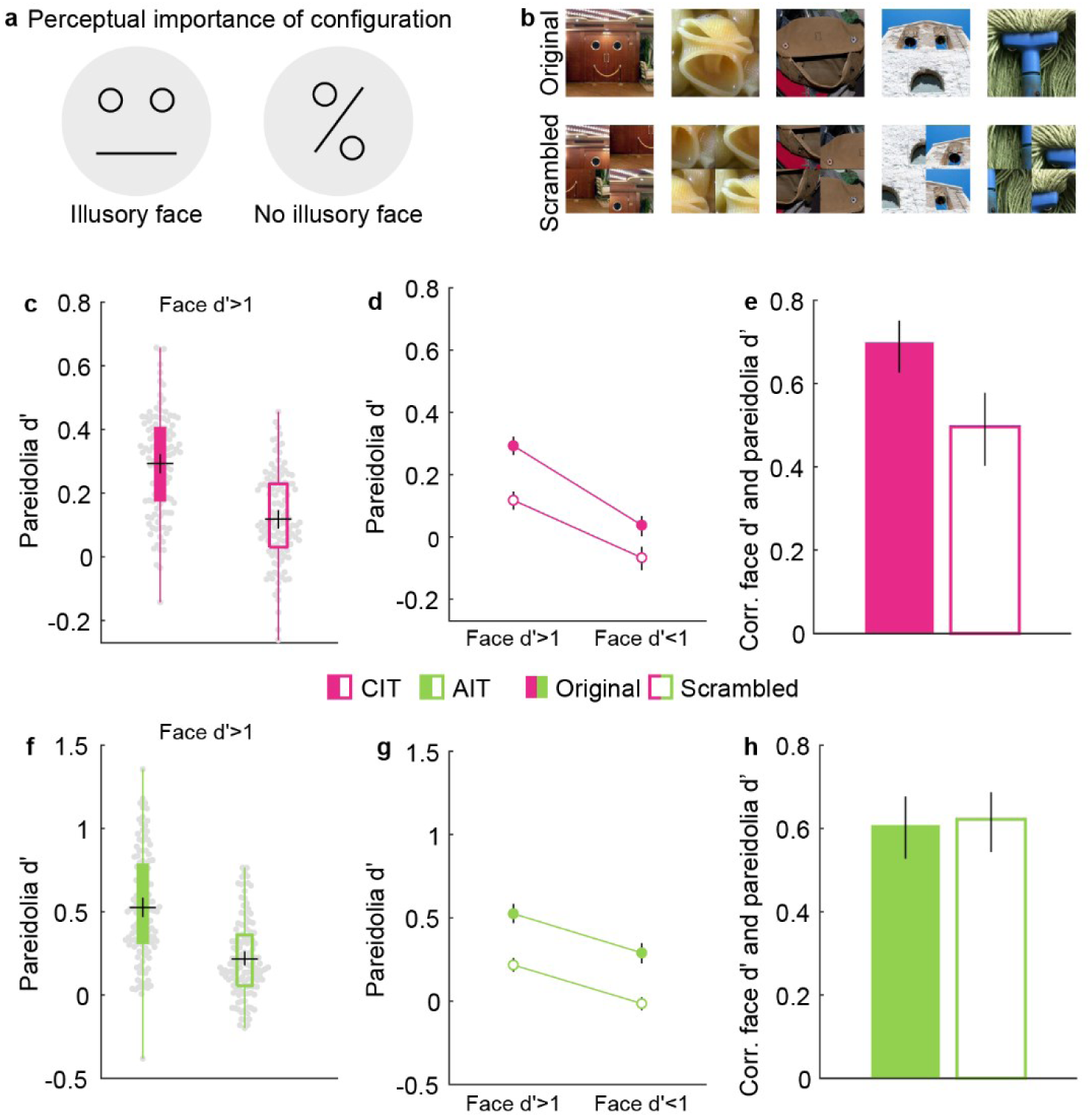
Quadrant scrambling reduces but does not eliminate the pareidolia selectivity. **a.** Cartoon example demonstrating the importance of configuration to invoke the perceptual experience of a face **b.** Example pareidolia images with their original (top) and quadrant scrambled versions (bottom). We divided each image into four equal quadrants and then shuffled the location of each quadrant, so that the individual features were maintained, but no longer in their original face-like configuration. **c.** Average pareidolia selectivity is shown in box plots for face units in CIT (113/208) for original (filled boxplot) and scrambled (open boxplot) images. The black central horizontal line shows the mean response, the black central vertical line indicates confidence intervals, and the bottom and top edges of the box depict the 25^th^ and 75^th^ percentiles, respectively. Whiskers (vertical lines in color) show the most extreme data points not considered outliers. Beeswarm plots behind the box plots show the pareidolia selectivity of each individual unit represented by a grey dot. **d**. Plot showing the average pareidolia selectivity of face (face d’>1; n = 113) and nonface (face d’<1; n = 95) units for original (filled) and scrambled (open) pareidolia images. Error bars indicate confidence intervals. **e**. Bar plots showing the correlation of face selectivity with pareidolia selectivity for original (filled bar) and scrambled (open bar) pareidolia images for all CIT units. Error bars indicate confidence intervals. **f.** Average pareidolia selectivity for face units in AIT (122/163). Same conventions as in **c**. **g.** Plot showing average pareidolia selectivity for face (face d>1; n = 122) and nonface (face d’<1; n = 41) units in AIT. Same conventions as in d. **h.** Bar plots showing correlation of face selectivity with pareidolia selectivity for all units in AIT. Same conventions as in **e**.

To address this question, we divided each image into four quadrants, and then shuffled the quadrants, so that the new image retained the individual features of the pareidolia image but lost the original face-like spatial configuration (Fig. 3b). Strikingly, we found that quadrant scrambling of the pareidolia images did not eliminate pareidolia selectivity of face units (CIT = 0.12 ± 0.16, t_112_ = 8.04, p = 1.05 x 10^-12^, AIT = 0.22 ± 0.23, t_121_ = 10.3, p = 3 x 10^-18^). Further, pareidolia selectivity of face units was still significantly higher than pareidolia selectivity of nonface units (CIT: t_206_ = 7.8, p = 3.75 x 10^-13^, AIT: t_161_ = 6.04, p = 1.02 x 10^-8^). The positive correlation between face and pareidolia selectivity was also preserved (CIT: n = 208, Pearson’s r = 0.5, p = 2.49 x 10^-14^, AIT: n = 163, Pearson’s r = 0.62, p = 7.95 x 10^-19^) and significant for each of the 8 monkeys (Table S1; CIT: t_3_ = 2.4, p = 0.049; AIT: t_4_ = 4.61, p = 0.005). Thus, quadrant scrambling the pareidolia images removed the illusory perception of a face but did not eliminate pareidolia selectivity or the association between face and pareidolia selectivity.

Nevertheless, scrambling the global configuration of the pareidolia images does cause a significant reduction in the average pareidolia selectivity of face cells. Thus, we asked if scrambling the global configuration also impacts nonface units. If some of the pareidolia selectivity of face cells is driven specifically by the facial configuration, the impact of scrambling should be restricted to face cells (i.e., neurons that could potentially respond selectively to a facial configuration). In Fig. 3c-h, we compare pareidolia selectivity and its relation to face selectivity for original vs scrambled images. In CIT, scrambling the global configuration significantly reduced the average pareidolia selectivity for face units (Fig. 3c, d; original = 0.29, scrambled = 0.12, t_112_ = 9.98, p = 3.55 x 10-^17^). However, scrambling the global configuration also significantly reduced the pareidolia selectivity of nonface units (Fig. 3d; original = 0.04, scrambled = - 0.07, t_94_ = 9.04, p = 2.02 x 10^-14^), although the effect of scrambling was significantly larger for the face units than for the nonface units (Fig. 3d, t_206_ = 3.23, p = 0.0015). Indeed, this is also reflected in the significant reduction by scrambling in the correlation between face selectivity and pareidolia selectivity across all CIT units (Fig. 3e; n = 208, original r = 0.7, scrambled r = 0.5, difference in r = −0.2, 95%CI [-0.28, −0.11]). These results suggest that scrambling the face-like configuration has a larger effect on the face units than the non-face units in CIT. In AIT, scrambling the pareidolia images also significantly reduced the average pareidolia selectivity for face units (Fig. 3f, g; original = 0.53, scrambled = 0.22, t_121_ = 16.5, p = 9.32 x 10^-33^). It also significantly reduced the average pareidolia selectivity for nonface units (Fig. 3g; original = 0.3, scrambled = −0.01, t_40_ = 13.7, p = 1.002 x 10^-16^). However, the effect of scrambling on face units was not different from the effect of scrambling on nonface units (Fig. 3g, t_161_ = 0.098, p = 0.92). Correspondingly, there was also no difference in the correlation between face and pareidolia selectivity across all AIT units (Fig. 3h; original r = 0.61, scrambled r = 0.62, difference in r = 0.01, 95%CI [-0.05, 0.08]). Thus, in AIT, the effect of scrambling on pareidolia selectivity was independent of face selectivity. We provide in the discussion a potential explanation as to why nonface units in CIT are less affected by scrambling, unlike in AIT. Overall, our results show that, though quadrant scrambling reduced the pareidolia selectivity in CIT and AIT, it did not eliminate it. Further, this reduction was largely independent of face selectivity, suggesting that scrambling the global configuration affects IT neurons’ responses generally, independently of their face selectivity and thus independent of their potential selectivity for a facial configuration.

Given that original and scrambled versions of the pareidolia images were presented together in the first experiment, the preserved pareidolia selectivity observed for the scrambled versions could be attributed to familiarity with the original images. To rule out this potential confound, we conducted a second experiment where the monkeys were presented with only scrambled versions of images for which they had never seen the original version. We found that even for unfamiliar scrambled images, face units in CIT and AIT were pareidolia selective (CIT = 0.1 ± 0.16, t_85_ = 5.79, p = 1.14 x 10^-7^, AIT = 0.14 ± 0.15, t_93_ = 8.78, p = 7.72 x 10^-14^), with a significantly positive correlation between face and pareidolia selectivity (Fig. S1a, b, CIT: n = 162, Pearson’s r = 0.28, p = 2.54 x 10^-4^, AIT: n = 114, r = 0.47, p = 1.43 x 10^-7^). Thus, scrambled-pareidolia selectivity in face units does not depend on familiarity of the monkeys with the original, unscrambled, pareidolia images.

### Pareidolia selectivity is primarily driven by “eye” quadrants

Although quadrant scrambling did not abolish the pareidolia selectivity of face cells, it did cause a reduction in pareidolia selectivity. Was this reduction because scrambling disrupts the spatial configuration of the object parts or because it changes the absolute position of individual parts? To answer this question, in a third experiment, we took the top 50 pareidolia images from Experiment 1 based on neural responses and their matched control objects, and presented the full original or scrambled versions, but also the four quadrants in isolation (Fig. 5a). Since quadrant scrambling introduces edges that are normally absent in the original images (see Fig. 3a for examples), in this experiment we also introduced a grey cross in both sets of full images. For clarity, when discussing the results of this experiment in this section, we report the average pareidolia d’ for the different image manipulations as a percentage of the average pareidolia d’ based on the full original image for face units only (face d’>1). Further, to compute the pareidolia d’ based on the individual quadrants, we used the individual quadrant of the pareidolia images and the corresponding quadrant of the matched control objects at the same location. Note that we compare pareidolia d’ rather than absolute response magnitudes, because presenting scrambled images or isolated quadrants (i.e., smaller images) can affect the overall response magnitude for all images, without necessarily affecting the pareidolia selectivity. Finally, we included only 34/50 pareidolia images in our analysis, where the eyes and mouth were clearly separable and confined to different quadrants.

We first assessed how the average pareidolia d’ of face units for the scrambled full images and for the sum of the isolated quadrants compared to the pareidolia d’ for the original images. In central IT (Fig. 4b), the average pareidolia selectivity based on scrambled full images was only 21% of the average for the original full images (t_68_ = 11.2, p = 4.63 x 10^-17^). This reduction in pareidolia selectivity could be because of a change in the absolute position of the object parts or because of a change in configuration. To isolate the effect of configuration from the effect of position, we looked at the average pareidolia d’ based on the sum of the responses to the individual four image quadrants (of pareidolia images and controls) presented in their original position. We found that the average pareidolia selectivity based on the summed response of these four quadrants in central IT was 78%, much higher compared to the 21% for the scrambled full images. Thus, removing only the global configuration (face-like or otherwise), while retaining the position of the object parts, reduced pareidolia selectivity by only 22% (100% - 78%; t_68_ = 3.7, p = 0.00046). Interestingly, the average pareidolia selectivity based on the sum of the response to isolated quadrants of the scrambled images, was 46%. Thus, changing the position of the object parts to the scrambled image locations further reduced pareidolia selectivity by 32% (78% - 46%; t_68_ = 5.13, p = 2.62 x 10^-6^). Further, the average pareidolia selectivity for individual quadrants in scrambled locations was still twice as high as the pareidolia selectivity for the full scrambled images (46% - 21%; t = −3.8, p = 2.68 x 10^-4^), suggesting that there is a *negative* effect of the scrambled configuration on pareidolia selectivity. Taken together, these results suggest that quadrant scrambling reduced pareidolia selectivity in central IT for three reasons: because it moved the parts to a new position, it removed the face-like configuration, and it introduced a new configuration.

**Fig. 4.**
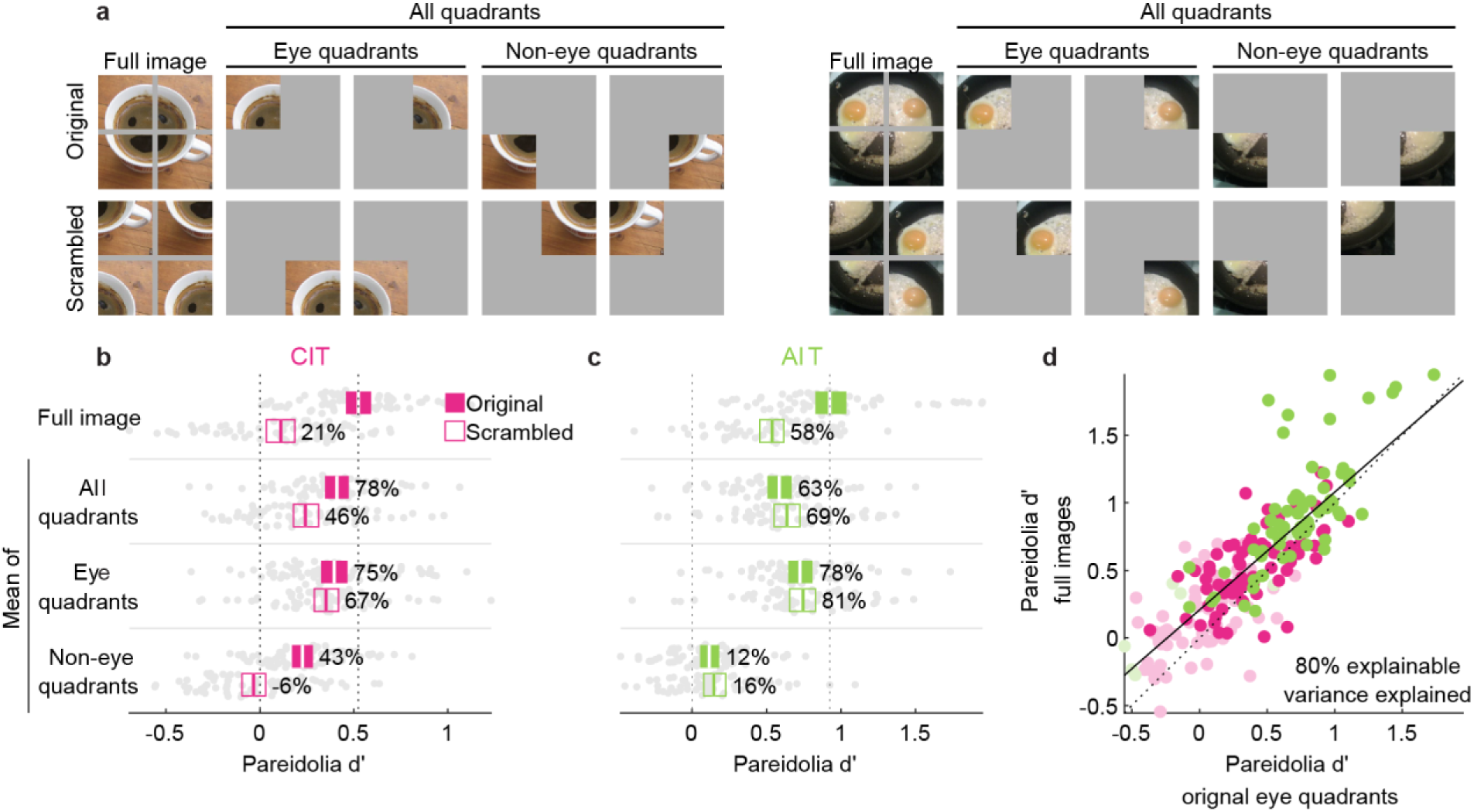
Pareidolia selectivity in face units is primarily driven by “eye” quadrants. **(a)** Two example pareidolia images, and the quadrants that were presented in isolation. A gray cross was introduced in the full original image to ensure the shared presence of quadrant edges in both original and scrambled images. The gray background in the single-quadrant images was the same color as the screen background, which made it appear as if the quadrants were presented in isolation. The quadrants were divided into “eye” and “non-eye” categories to investigate their contribution to pareidolia selectivity of face-cells. **(b)** The pareidolia d’ in CIT for face units (face d’>1; n = 69) for full images, the average d’ for the sum of all quadrants, the average d’ for the sum of only the eye quadrants and the average d’ for the sum of only the non-eye quadrants for original (filled) and scrambled (open) images. Beeswarm charts show the d’ of each neural unit, where each grey circle represents a neural unit. Colored boxes indicate 95% bootstrapped CI, calculated by resampling sites, with the central vertical line indicating the mean d’ across units. The numbers adjacent to the colored boxes indicate the percentage of the mean d’ value relative to the d’ of the full original images. **(c).** Same conventions as in (b), except for face units in AIT (n = 65). **(d)** Scatter plot showing distribution of pareidolia d’ for the eye quadrants only from original images on the x-axis, and the pareidolia d’ for the full original image on the y-axis. Each circle represents a neural unit in CIT (pink) and AIT (green). Lighter circles are face cells with face d’<1 and darker circles represent units with face d’>1.

Next, we categorized the quadrants into “eye” and “non-eye” quadrants, depending on which part of the illusory face they represented in the original pareidolia image, to investigate their individual contribution to pareidolia selectivity. We found that the average pareidolia selectivity for the sum of the eye quadrants was 75% of the average for the full original images, which is not significantly different from the selectivity based on the sum of all quadrants (78% - 75%; t_68_ = 0.75, p = 0.46). Furthermore, pareidolia selectivity based on the sum of individual eye quadrants in the scrambled location was 67%, which is not significantly different from the selectivity for the sum of individual eye quadrants in their original location (75% - 67%; t_68_ = 1.4, p = 0.18). Finally, there was still significant pareidolia selectivity for the sum of non-eye quadrants in the original location (43%), but not in the scrambled configuration (−6%). Overall, these results suggest that pareidolia selectivity in CIT is driven mostly by eye parts, irrespective of their position, and reduced by the scrambled positions of the non-eye parts.

In anterior IT (Fig. 4c), scrambling the configuration significantly reduced the average pareidolia selectivity (t_64_ = 9.92, p = 1.46 x 10^-14^) to 58% of the average for the full original images, suggesting that the impact of scrambling is less profound than in central IT. This value was comparable to the average pareidolia selectivity values based on the sum of the four quadrants in the original positions (63%). Thus, removing only the global configuration while retaining the position of each quadrant, reduced pareidolia selectivity by 37% (100% - 63%; t_68_ = 3.7, p = 0.00046). Further, there was no major negative effect of the scrambled configuration, i.e., when the quadrants took on new positions (63% - 58%; t = 1.7, p = 0.095). Consistent with this, the pareidolia selectivity based on the sum of quadrants in the scrambled positions was comparable (69%), suggesting that the absolute position of the scrambled quadrants did not reduce pareidolia selectivity. Interestingly, the average pareidolia selectivity based on only the sum of eye quadrants was even closer to the average for the full original images (78% and 81% for original and scrambled positions, respectively). Moreover, pareidolia selectivity for the non-eye quadrants was negligible (12-15%) compared to the magnitude for the full original images. Taken together, our results suggest that—like central IT—pareidolia selectivity in anterior IT is largely explained by eye quadrants, with a relatively smaller contribution of the face-like configuration. Indeed, this result can also be observed in Fig. 4d, where pareidolia selectivity based on eye quadrants explains 80% of the variance explained for the pareidolia selectivity of the full original images. However, unlike central IT, for anterior IT we found no major additional negative effect of the scrambled configuration or quadrant positions per se on pareidolia selectivity.

Overall, these results indicate that pareidolia selectivity is primarily attributable to the pareidolic eye features, i.e., individual object parts that, when arranged in the right configuration, act as eyes of the illusory face. The face-like configuration itself appears to contribute minimally to pareidolia selectivity, much less than simply comparing full scrambled and original images might suggest. In fact, particularly in the case of CIT, our results imply a negative effect of the absolute and relative positions of “non-eye” parts in scrambled images that is larger than the positive effect of the face-like configuration in original images.

### Pareidolia selectivity can be captured by a non-face encoding model

Our results thus far suggest that pareidolia selectivity of face cells is primarily driven by individual object parts or local features and does not require an arrangement in a face-like configuration. This suggests that neural pareidolia selectivity largely results from a tuning for more generic features that apply to all kinds of objects, even ones that do not resemble a face. That is, certain generic features, like the dark holes in a leather belt or the roundness of an egg yolk, are likely more prevalent in images with illusory faces, but can also be present in the matched control objects. Does neural tuning for such individual features also predict pareidolia selectivity?

In recent work, we showed that face selectivity can be predicted from features that apply to non-face objects(Vinken et al., 2023). Applying the same approach from our previous work, we quantified the tuning of each neural site based on the characteristics of images without illusory or real faces. We then assessed if this tuning could predict pareidolia selectivity in face cells. We computed the neural tuning by fitting an encoding model based on a convolutional neural network (CNN) trained on object categorization (Krizhevsky et al., 2012). The model was pretrained on ImageNet, which does contain some faces, albeit not in a separate category. Critically, we used only matched controls (of the pareidolia images) or non-face object images to linearly map (fit) the features encoded by the CNN to the neural responses (see Methods and Fig. 5a). We term this the non-pareidolia encoding model, because it characterizes the neural tuning using only responses to images that are not faces and do not contain illusory faces. Because an encoding model can be derived only from reliable responses, we excluded 105 sites (face d′ between - 1.9 and 5.4) that had response reliability below the threshold of 0.4 for matched controls, leaving 266 remaining sites (face d′ between −2.3 and 6.1). This exclusion did not qualitatively affect the results. We then used the model-predicted responses to the pareidolia images and matched controls to compute a model-predicted pareidolia-selectivity value for each neural site. We found that the non-pareidolia encoding model accurately predicted the observed pareidolia selectivity (Fig. 5b; CIT: Pearson’s r between observed and predicted values = 0.71, p = 1.1 x 10^-27^, AIT: Pearson’s r = 0.76, p = 2.5 x 10^-19^). Furthermore, this model-predicted pareidolia selectivity correlated with the face selectivity observed in the actual neural responses (Fig. 5c; CIT: Pearson’s r = 0.68, p = 4.3 x 10^-24^, AIT: Pearson’s r = 0.56, p = 4.5 x 10^-9^). Hence, even though no pareidolia or face images were involved in quantifying the feature-tuning with the encoding model, the model successfully predicted both pareidolia selectivity and its relationship to the face selectivity of the actual neural sites.

**Fig. 5.**
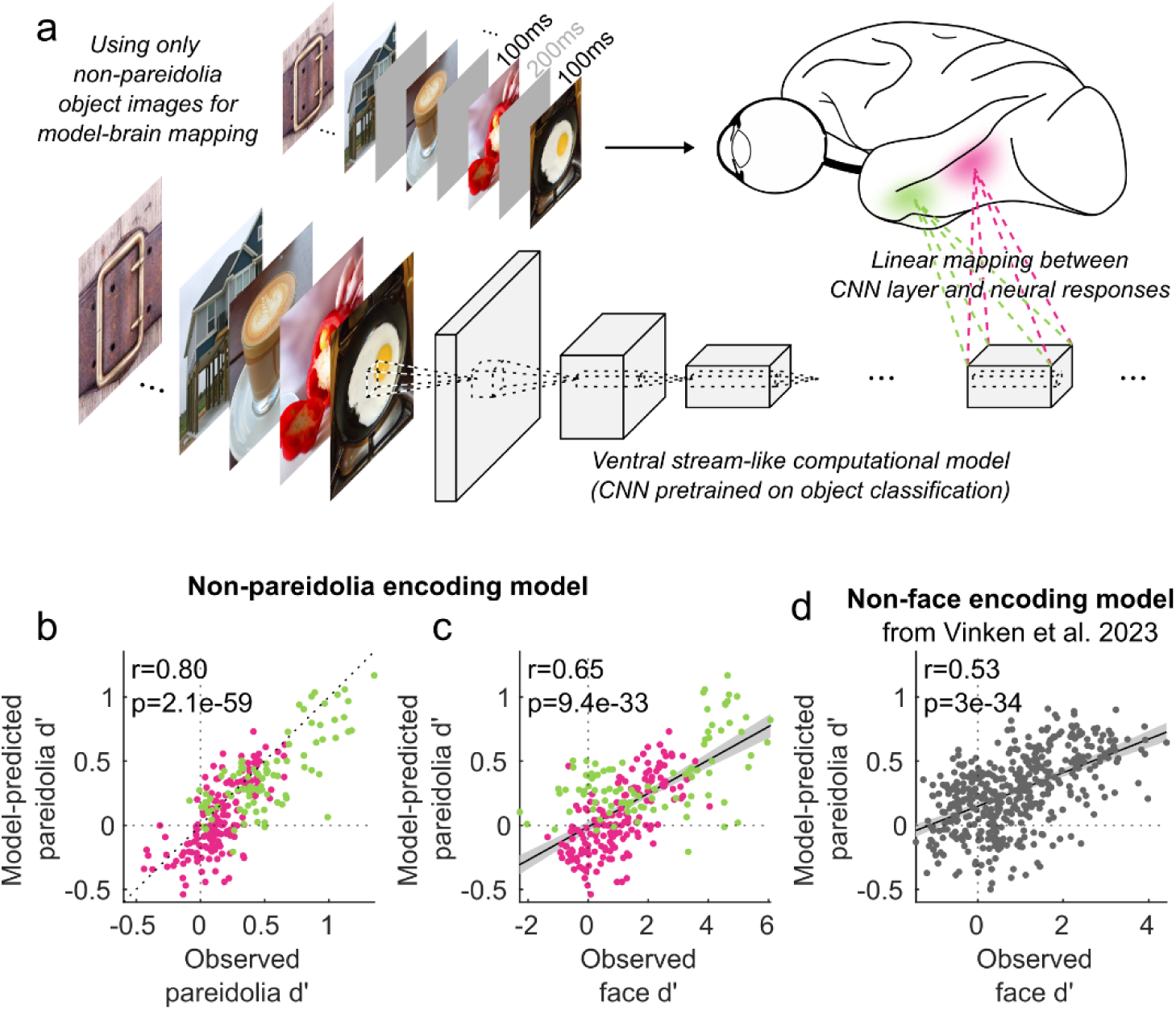
Pareidolia selectivity is predicted by the feature-tuning estimated from non-pareidolia, non-face images. **a.** Design of the non-pareidolia encoding model. This encoding model was based on a CNN trained on object classification. We input into this model the same images that were also presented to the monkeys. Using cross-validated subsets of only the 100 matched controls and 40 non-face objects, we estimated a linear mapping from the model to each neural site (see Methods). This resulted in an encoding model for each neural site that captures the tuning for characteristics of only non-pareidolia, non-face images. **b.** Scatterplot showing the similarity between each neural site’s observed (i.e., computed from neural responses) pareidolia d’ and its corresponding model-predicted value. Each dot represents a neural site in CIT (pink; n=171) or AIT (green; n=95). **c.** Scatterplot showing the correlation between neural face selectivity (observed face d’) and model-predicted pareidolia selectivity (pareidolia d’). Same conventions as b. **d.** Scatterplot showing the correlation between observed neural face selectivity and model-predicted pareidolia selectivity, using the independent non-face encoding model from Vinken and colleagues (Vinken et al., 2023)

Finally, as a more stringent test of our hypothesis that generic object features predict pareidolia selectivity, we input both pareidolia and control images into the non-face encoding model of (Vinken et al., 2023). Briefly, this independent model was constructed using a CNN-based encoding model (using the same base model as above) and was fit (linearly mapped) on the responses from 449 central IT sites to 932 exclusively inanimate, non-face objects (for more details see Vinken et al., 2023). No actual pareidolia images or matched controls were shown in this experiment. Importantly, this independent non-face encoding model, which stems from a different study with entirely separate data and experimental protocols, could predict a degree of pareidolia selectivity that positively correlated with the face selectivity observed in the neural sites upon which the model was originally based (Fig. 5d; Pearson’s r = 0.53, p = 3 x 10^-34^). Thus, the link between pareidolia selectivity and face selectivity is captured by domain-general features, that apply to non-pareidolia, non-face images and are derived from a rich feature space that was trained to represent all kinds of objects.

### Faceness does not explain pareidolia selectivity in face cells

Assuming that a global, face-like configuration is critical for the perceptual experience of a face, our results indicate a discrepancy between the cues for the human perception of faceness and those that activate face cells. One way to explore this dissociation between visual perception and neural response magnitude, is to investigate how the neural response of face cells to pareidolia images relate to human “faceness” ratings of the same images. To this end, in a fourth experiment, we presented the monkeys with 200 pareidolia images and their matched controls, that human subjects had rated on an 11-point scale from 0 “cannot see a face” to 10 “easily see a face” in a recently published study (Wardle et al., 2022). If face-cell responses to illusory faces encode how much a stimulus looks like a face, then face-cell responses should be equal for controls and pareidolia images that are matched in faceness. To test this, we selected an image subset consisting of the 10 least face-like pareidolia images (faceness rating: 2.9 ± 0.8) and the 10 most face-like control images (faceness rating: 3 ± 0.5; Figure 6a). Then we computed the pareidolia d’ using the selected subset of 10 pareidolia images and 10 control objects, which were not necessarily matched in identity. We found that, even though these 10 pareidolia and 10 control images were, on average, matched in faceness ratings, face units responded more to illusory faces than to controls (pareidolia d’, CIT: 0.16, t_86_ = 3.8, p = 0.0002; AIT: 0.15, t_119_ = 4.1, p = 8 x 10^-5^; Figure 6b). Thus, pareidolia selectivity persists even when there is no difference in faceness ratings.

**Fig. 6.**
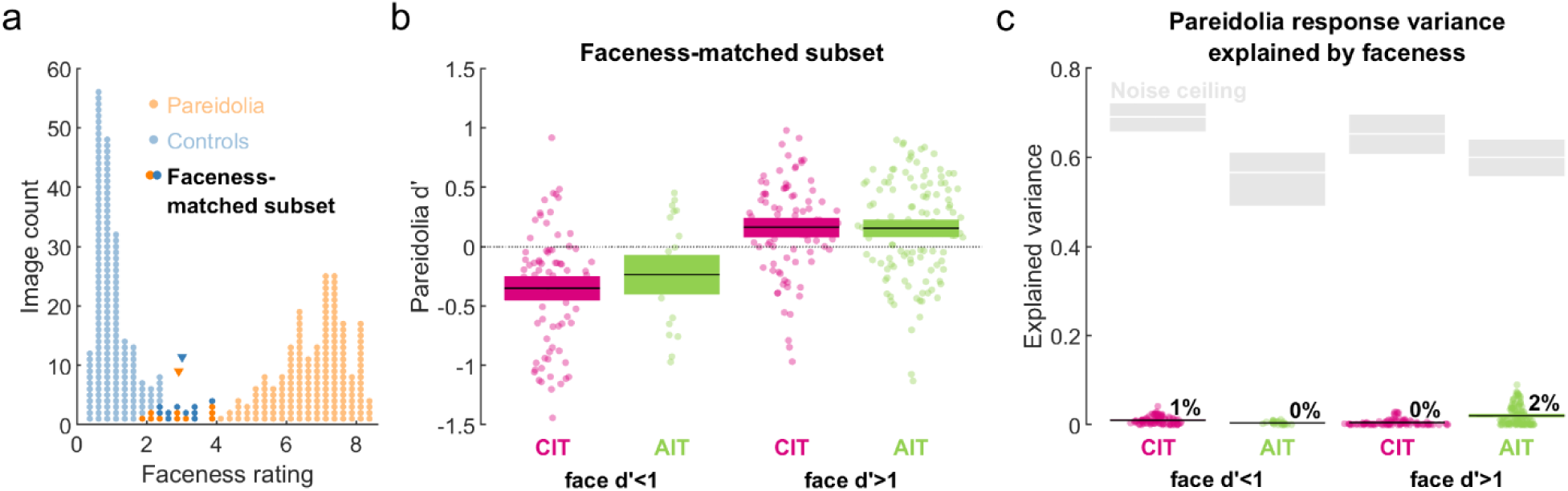
Faceness does not explain pareidolia selectivity in face cells. (a) Distribution of perceptual faceness ratings for the 200 pareidolia images (blue dots) and their matched controls (red dots) taken from Wardle et al, 2022. Darkened dots indicate a faceness-matched subset of the 10 most face-like control images and the 10 least face-like pareidolia images. Downward-pointing triangles indicate the means of the faceness-matched subsets. (b) Pareidolia d’ values for the faceness-matched subset. Each marker represents a single neural site. Colored boxes: 95% bootstrapped CI, calculated by resampling sites; black line: mean. (c) Response variance for pareidolia images explained by faceness ratings. Black line: mean. Colored boxes indicating 95% bootstrapped CI are obscured by the black mean line. Grey boxes and the white line indicate the 95% bootstrapped CI and mean across neural sites of the noise ceiling (i.e., the maximum explainable variance, computed as the neural response reliability multiplied by the reliability of the faceness-rating).

Next, to further investigate the relation between neural site responses and faceness ratings, we investigated whether the response differences between individual pareidolia images could be explained by differences in faceness. For this analysis, we used all 200 pareidolia images. We computed for each unit the variance in pareidolia responses explained by faceness ratings (squared Pearson correlation between responses and faceness ratings). For face units, the faceness ratings explained on average ∼0% (95%CI[0.3 0.6]) of the pareidolia response variance in CIT, and ∼2% (95%CI[1.7 2.5]) in AIT (Figure 6c). For units with face d’<1, the explained variance was ∼1% (95%CI[0.8 1.2]) in CIT and ∼0% (95%CI[0.3 0.6]) in AIT. Thus, faceness ratings did not explain any notable response variance for pareidolia images. Even if responses of individual neural sites do not encode the perceptual faceness of pareidolia images, it could still be that the perceptual faceness can be linearly decoded from the neural population. To test this, we fit a cross-validated linear regression (see Methods) predicting faceness ratings from the population response pattern. For both CIT and AIT, there was a weak correlation between observed and predicted faceness (CIT: r_198_=0.16, p=0.03, 95%CI[0.02 0.29]; AIT: r_198_=0.26, p=0.0002, 95%CI[0.12 0.38]). The square of these correlations is 2.5% for CIT (noise ceiling 61.4%) and 6.7% for AIT (noise ceiling 78.1%), which is only slightly higher than the average explained variance for individual neural sites (∼0-2%). Taken together, these findings further support the notion that pareidolia selectivity of face-cells is not driven by the perceived “faceness” of the pareidolia images.

## Discussion

In this study, we used face pareidolia, which refers to seeing illusory faces in inanimate objects, to study holistic processing in face cells. Our aim was to test whether face selective units in the IT cortex respond to illusory faces because of the face-like configuration of individual object features, which is integral to the perceptual experience of a face. Overall, we found that face selective units (i.e., units that respond more strongly to faces than nonfaces) in macaque central and anterior IT cortex were also highly pareidolia selective (i.e., they responded more strongly to pareidolia images than to matched control objects). When investigating what drives pareidolia selectivity in face cells, we found that 1) it did not require the global configuration that is critical for the perceptual experience of an illusory face, 2) it was instead primarily driven by local features, which explained most of the selectivity even when presented in isolation, 3) it could be explained by generic features also present in non-pareidolia nonface objects, and 4) it could not be explained by human faceness ratings. Taken together, our results suggest that neural pareidolia selectivity of macaque face cells is dissociable from the extent to which an object looks like a face. Instead, it can be explained mostly by isolated pareidolic eye features that that merely represent object parts without the context of a face-like configuration.

One interesting result was that face selectivity was negatively correlated with object rank correlation (a metric we used to assess if neural responses were selective to features shared between pareidolia images and their matched control objects), particularly in anterior IT. Given that pareidolia images and matched controls share features associated with object identity (such as shape, color, or texture), this result suggests that something other than these shared features drives the face-cell responses, possibly the presence of the face-like configuration (Decramer et al., 2021). If this is the case, then removing that configuration should impact only neurons that encode a face-like configuration and thus, are face selective. On the contrary, we found that scrambling the configuration caused a general reduction in *all* units in CIT and AIT and was not restricted to face cells. We do note that the reduction caused by scrambling was smaller for the nonface units than for the face units in CIT. A potential explanation is that, compared to AIT, the pareidolia selectivity for the original images of nonface units in CIT was lower (CIT: 0.04, AIT: 0.3) and therefore there was less range for those values to reduce as an effect of scrambling. Overall, the fact that scrambling equally (AIT) or almost equally (CIT) affected nonface cells suggests that pareidolia selectivity in face cells is not explained solely by the face-like configuration of the object features. Moreover, it is also important to note that face cells remained pareidolia selective (albeit significantly less) even when the global configuration of the pareidolia images was scrambled. This suggests a significant contribution of local features in driving face cell responses, which are not affected by scrambling. In this way, face cells appear no different from other category-selective IT neurons, which have been shown to encode local features or object parts, rather than a global spatial arrangement of object features (Ayzenberg & Behrmann, 2022; Grill-Spector et al., 1998; Guo et al., 2022; Jagadeesh & Gardner, 2022; Vogels, 1999).

The downside of using configuration scrambling as a method to investigate holistic processing is that it does not consider if any change in response was caused by breaking up the configuration or because it changed the absolute position of certain object parts (Jagadeesh & Gardner, 2022). An alternative is to present isolated features versus combinations thereof. Using this approach, one study suggested that face cells needed combinations of features to drive their responses, whereas others found that face cells could be driven by isolated features as well (Freiwald et al., 2009; Issa & DiCarlo, 2012; Waidmann et al., 2022). Here, we confirm, using both scrambled images and isolated parts, that face-cell responses did not require a holistic, face-like configuration. Moreover, we found that the pareidolia selectivity could be mostly explained (80%) by the pareidolic eye features alone. While the significance of eyes for face cells have previously been demonstrated (Azadi et al., 2023; Freiwald et al., 2009; Issa & DiCarlo, 2012; Waidmann et al., 2022), an important distinction to be made here is that real eyes are still recognizable as eyes, even without the context of a real face. On the other hand, when viewed in isolation, we argue that a pareidolic eye feature (e.g., an egg yolk in a frying pan) does not resemble an eye more so than it resembles the actual object part (i.e., an egg yolk). Our findings also suggest that face cells do not require actual face parts to give selective responses, and that local object features, or object combinations are enough to drive that selectivity, even when the face is occluded or entirely absent (Arcaro, Ponce, et al., 2020; Meyers et al., 2015; Vinken et al., 2023). Thus, our results suggest a tuning that is largely driven by low level features such as contrast or shape, as opposed to a tuning driven by a face-like configuration. Such low-level features are present in object parts as well as in actual eyes and possibly contribute to the detection of a face (Taubert et al., 2022). Some evidence for this has also been observed in causal studies where the effect of stimulating face and body-selective IT neurons is not limited to the preferred categories, but also affects the perception of other categories, such as non-faces for face cells and faces and houses for body cells(Kumar et al., 2022; Moeller et al., 2017). Overall, our results provide further support for the notion that, rather than encoding high-level, category-specific features, single neurons in IT cortex owe their category selectivity to learnt combinations of lower-level, domain-general features that together form an integrated object space (Arcaro & Livingstone, 2017; Bao et al., 2020; Doshi & Konkle, 2023; Konkle & Caramazza, 2013; Vinken et al., 2023).

Our results show that 75-80% of the variance and magnitude in pareidolia selectivity was explained by eye quadrants only. This leaves open the question of what other factors contributed to the relatively smaller 20-25% of pareidolia selectivity left unexplained. One possibility is that there was a significant contribution of the face-like configuration in pareidolia images. This could also be the arrangement of smaller subsets of parts, such as two pareidolic eye features, and not the whole face-like configuration per se (Freiwald et al., 2009). However, several of our results instead point to a substantial contribution of characteristics that are not face-like. First, configuration scrambling reduced selectivity not just in face cells, but in *all* units, suggesting a contribution of features that are not correlated with faces. These features could be nonface-like configurations of object parts, as well as larger properties of the whole object upon which the face-like arrangement is embedded (e.g., the large round frying pan, fruit, etc.). Second, the result that pareidolia selectivity was lowest for scrambled images (compared to summing across all the individual quadrants) points to a negative effect of the spurious configuration introduced by scrambling, particularly in CIT. This suggests that face cells are at least modulated by nonface-like configurations. Thus, further research will be necessary to disentangle the potential contribution of face-like configurations in pareidolia selectivity from other nonface-like characteristics hinted at by our results. Broadly, our results indicate that face cells largely encode local object features, as opposed to global configurations, like other category-selective neurons in IT. This suggests that the neural mechanisms underlying face processing are not different from general object processing(Hesse & Tsao, 2020; Tan & Poggio, 2016). It remains an open question where the holistic processing for illusory faces occurs, since, perceptually, we do perceive faces in inanimate objects in the original but not scrambled configuration. Under the assumption that face perception culminates in IT, our results suggest that the holistic perception of illusory faces could arise from a largely featural cortical representation (Jagadeesh & Gardner, 2022). That is, a holistic perceptual experience could emerge from configural information that is distributed across many neurons, which would explain the relatively small configural effects we found in individual units. Another possibility is that the integration of features underlying perceptual holistic processing might instead occur in regions downstream from IT cortex (Bonnen et al., 2021; Waidmann et al., 2022).

There are a few limitations to our study. First, while humans clearly perceive illusory faces in pareidolia images, even assigning age, gender, and emotion to these nonface objects(Wardle et al., 2020, 2022), it is unclear whether monkeys even perceive illusory faces. Eye-tracking experiments show that free-viewing monkeys look preferentially at pareidolia images, and their fixation patterns for pareidolia images were correlated with their viewing patterns for faces (Azadi et al., 2023; Taubert et al., 2017). However, the monkeys also fixated small round object parts regardless of the presence of an illusory face. Thus, a behavioral preference for illusory faces could simply be a preference for small round shapes, which are more prevalent in illusory faces, without necessarily being an indication that the monkeys perceive illusory faces. Second, we did not quantitatively assess whether the global configuration of pareidolia images is essential for a human perceptual experience of a face in the pareidolia images used in our study. Future experiments in humans could test if pareidolia images with 2 x 2 scrambling, like we did here, would still be perceived as face-like and if face-selective regions in humans are also robust to scrambling the spatial configuration of pareidolia images. Nevertheless, it is important to highlight that the phenomenon of face pareidolia occurs specifically for visual patterns with a spurious face-like arrangement, which can be separated from the more general faceness of an object. Any object, such as an apple or a house, could be evaluated on how face-like it is (Bardon et al., 2022), without necessarily creating the perceptual illusion of a face. Finally, our inclusion of the face-deprived monkeys in the current study might seem counterintuitive to the hypothesis we tested. Previous work from our lab has shown that monkeys reared without experiencing faces for the first year of their life do not develop face-selective domains in the IT cortex(Arcaro et al., 2017). However, we decided to include them since both monkeys had more than two years of experience with faces when we ran these experiments and fell within the range of face d’ demonstrated by the control monkeys (Fig. S2, t_7_ = 0.08, p = 0.94). Importantly, excluding them from our analysis did not qualitatively change any of our results.

In conclusion, we show that face cells in IT cortex are pareidolia selective and that face selectivity is correlated with pareidolia selectivity. Surprisingly, scrambling the face-like configuration of the face pareidolia images did not abolish the pareidolia selectivity of face cells. Instead, pareidolia selectivity was primarily driven by pareidolic eye features, which represent object parts or object features when viewed in isolation. Moreover, the pareidolia selectivity of face cells could be captured by an encoding model fit on non-face, non-pareidolia images. Finally, we found that human faceness ratings did not explain any notable response variance of face units. Taken together, our results suggest that face cells encode local visual features, without requiring the spatial arrangement of the features, contrary to visual perception in primates, which requires holistic information (Ayzenberg & Behrmann, 2022; Guo et al., 2022; Jagadeesh & Gardner, 2022; Parr, 2011; Taubert et al., 2012). This suggests that face cells encode object features similarly to other category selective neurons in IT (Hesse & Tsao, 2020; Tan & Poggio, 2016) and provides further evidence for the notion that face cell activity is not restricted to only faces but is instead indicative of a broader tuning in an integrated object space (Bao et al., 2020; Decramer et al., 2021; Doshi & Konkle, 2023; Taubert et al., 2020; Vinken et al., 2023). More generally, our results underscore the importance of exercising caution when interpreting neural activity, as it may not always mirror our conscious perceptual experiences. It is tempting to understand selective responses in terms of what we, as human observers, see. However, in doing so, one might end up imposing the biases of human perception itself, such as seeing a face where none exists, onto neural activity.

## Methods

All procedures were approved by the Harvard Medical School Institutional Animal Care and Use Committee and conformed to NIH guidelines provided in the Guide for the Care and Use of Laboratory Animals.

### Subjects and array location

Eight adult male macaques implanted with floating microelectrode arrays (32 channels, MicroProbes, Gaithersburg, MD or 128 channels, NeuroNexus, Ann Arbor, MI) or microwire bundles (64 channels; MicroProbes) were used in this experiment. The arrays were chronically implanted based on fMRI localization or anatomical “bumps”(Arcaro, Mautz, et al., 2020) in the lower bank of the superior temporal sulcus, at the location of the middle face region (ML/MF) for four monkeys and at the location of anterior face region (AL/AF) for five monkeys (one monkey had arrays in both middle and anterior face region). Two of the monkeys used in this experiment (one with an array in central IT, and one with arrays in both central and anterior IT) were face-deprived, i.e., they were prevented from seeing faces for the first year of their life (similar procedure, but different monkeys than the ones used in Arcaro et al. 2017(Arcaro et al., 2017)). However, at the time that these experiments were run, both monkeys had been exposed to faces for more than 2 years.

### Visual experiments

The monkeys sat upright in a plastic monkey chair, facing an LCD display screen 51 cm in front of the monkey. They were given continuous juice reward for maintaining fixation on a red dot (0.2 x 0.2°) in the center of the screen. Eye movements were tracked at 120 Hz using ISCAN system (Woburn, MA, http://www.iscaninc.com/). After array implantations, the receptive field of each array was mapped by presenting faces and fractals (5 by 5°) at 26 different locations on the screen relative to the fixation point. The images were presented at the rate of 100ms on, 200ms off, while the monkeys fixated the red dot in the center of the screen. For the main experiments, we used a rsvp-style task paradigm, where images (5 by 5°) were presented on a grey background at the rate of 100ms on, 200ms off at the center of the array’s mapped receptive field.

### Stimuli

The 100 pareidolia images used in the first experiment, each with their matched controls were obtained from Dr Jessica Taubert (some used in (Taubert et al., 2017; Wardle et al., 2020, 2022)). We also presented quadrant scrambled versions of the pareidolia and matched control images. To do this, we divided the image into four equal quadrants and then randomly shuffled the location of the quadrants (Fig. 3a). In the second experiment, to investigate the effect of familiarity, we presented only the quadrant scrambled versions of 119 pareidolia images and their matched controls that the monkeys had never seen before taken from Wardle et al, 2022(Wardle et al., 2022). In the third experiment, we took the top 50 images based on the neural response from Experiment 1 and presented the original and scrambled versions of the full images. To ensure the shared presence of edges in both original and scrambled images, we introduced a grey cross in both sets of images. To ensure that the grey cross did not obscure any part of the full image, the quadrants were moved slightly away from each other in the full images. We also presented the four quadrants in isolation for both original and scrambled images (see Fig. 5a for examples). The grey background in the quadrant only images was the same color as the screen background, which made it appear as if the quadrants were presented in isolation. This was done to ensure that the size of the quadrants was the same as in the original full images. Finally, in the fourth experiment, to compare the neural responses with the human faceness ratings, we presented the monkeys with all 200 pareidolia images and their matched controls from Wardle et al. (Wardle et al., 2022; the images they used in their experiment 2A; images and data available at https://osf.io/f74xh/). Each experiment also included 40 faces (20 human faces, 20 monkey faces) and 40 non-faces (20 familiar, 20 nonfamiliar) on a white background (Fig. 1a), which we used to quantify the selectivity of the neural units. The 20 non-familiar images of nonfaces are from Konkle et al. (Konkle et al., 2010), and were also presented in Ponce et al. (Ponce et al., 2019), while the 20 images of nonfaces familiar to the monkeys, human and monkey face images were taken in our lab.

### Non-pareidolia encoding model

The methods used for the CNN encoding model were largely the same as those in (Vinken et al., 2023). Briefly, we used an ImageNet-trained AlexNet model (Krizhevsky et al., 2012) as a base. We then fit a linear mapping between the activations of a chosen model layer and the trial-averaged neural responses for each neural site. We used the same model layer for all neural sites, by selecting the one that on average most effectively predicted responses to non-pareidolia images. For this linear mapping, we only used matched controls (n = 100) or non-face object (n = 40) images, thus excluding the pareidolia image (n = 100). This resulted in a non-pareidolia encoding model that quantifies neural tuning exclusively based on features present in non-pareidolia images.

We fit the linear mapping between model activations and neural responses as follows: First, we normalized the CNN layer’s outputs per channel using the SD and mean across all pareidolia, matched control, and non-face object images (and across locations for convolutional layers). We then reduced the dimensionality through principal component analysis. Next, we fit a linear support vector regression model, using 35-fold cross-validation, to map neural responses onto the principal components of the normalized CNN activations (using the MATLAB 2020a function ‘fitrlinear’, with the SpaRSA solver and regularization parameter lambda set to 0.01). Before fitting, we centered the predictors on the mean of the training fold, and both centered and standardized neural responses using the mean and SD of the training fold. We then evaluated performance based on all concatenated out-of-fold predicted responses.

For pareidolia images, which were excluded from the training folds, we calculated the image-wise average across out-of-fold predictions. Finally, we determined predicted pareidolia selectivity by calculating the pareidolia d’ using these out-of-fold predicted responses.

### Data analysis

#### Firing rates

Neural signals were amplified and sampled at 40 kHz using a data acquisition system (OmniPlex, Plexon, Dallas, TX). Multi-unit spiking activity was detected using a threshold-crossing criterion. The neural response was defined as the spike rate in a 150ms time window starting at 30-100ms after the image onset, depending on the monkey. The firing rates per neural site were trial averaged per image.

#### Response reliability

We determined the firing-rate reliability per neural site, which was used as a selection criterion for inclusion of neural units in further analysis. To calculate the reliability of the units, the number of repeated presentations or trials for each image were randomly split in half. Next, we created two response vectors, one per half of the trials, by trial averaging the neural response and computed the correlation between these two split-half response vectors. This step was repeated 100 times, each time using different random splits, and an average correlation *r* was computed by taking the mean of the 100 correlation values. The reliability *ρ* was computed by applying the Spearman-Brown correction the average correlation *r*:

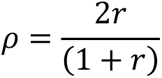

Only neural units with a split-half reliability *ρ* >0.4 were included in further analysis. For experiment 1, this resulted in the inclusion of 64/64 units from M1, 73/128 units from M2, 31/64 units from M3 and 40/64 units from M4, leading to a total of 208 recorded units in central IT. For anterior IT, we included 60/64 from M5, 13/64 units from M6, 24/64 units from M7, 57/64 units from M8 and 9/64 from M4, totaling 163 neural units. For experiment 2, we included 58/64 units from M1, 50/128 units from M2, 20/64 units from M3 and 34/64 units from M4, leading to a total of 162 recorded units in central IT and 49/64 from M5, 18/64 units from M7, 45/64 units from M8 and 2/64 from M4, totaling 114 neural units in anterior IT. For experiment 3, we included 58/64 units from M1, 56/128 units from M2, 23/64 units from M3 and 34/64 units from M4, leading to a total of 171 recorded units in central IT and 64/64 from M5, 4/64 units from M6, 19/64 units from M7, 54/64 units from M8 and 3/64 from M4, totaling 144 neural units in anterior IT. Finally, for experiment 4, we included 41/64 units from M1, 52/128 units from M2, 27/64 units from M3 and 25/64 units from M4, leading to a total of 145 recorded units in central IT and 10/64 from M5, 12/64 units from M6, 1/64 units from M7, and 49/64 units from M8, totaling 72 neural units in anterior IT.

For computing the 95% bootstrapped CI and mean across neural sites of the noise ceiling (depicted by grey boxes and white line in Fig. 6c), we used the neural response reliability multiplied by the reliability of the faceness-rating. In this case, the neural response reliability was computed as the rectified Pearson correlation between odd and even trial-averaged responses, with a Spearman-Brown correction (2r/(1+r)). For the faceness ratings, reliability was computed as the Pearson correlation between odd and even subject-averaged faceness ratings, with a Spearman-Brown correction.

#### Face selectivity

We quantified face selectivity using the d’ index, which compared trial-averaged responses to faces and to non-faces:

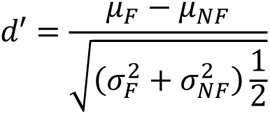

where *μ*_*F*_ and *μ*_*NF*_ are the across-stimulus averages of the trial-averaged responses to faces and non-faces, and *σ*_*F*_ and *σ*_*NF*_ are the across-stimulus standard deviations. We also used the same d’ metric to quantify pareidolia selectivity, which was computed based on neural responses to pareidolia images and matched controls.

#### Statistical inference

P-values were calculated using paired t-tests when comparing differences between conditions within face units, and unpaired t-tests when comparing face and nonface units. For R^2^ and correlations, which calculate the correspondence between two variables, permutation testing was performed by randomly shuffling one of the two variables. For the paired difference between two correlations, the condition labels were randomly shuffled for each pair of observations. 95% Confidence intervals were calculated using the bias corrected accelerated bootstrap(DiCiccio & Efron, 1996), based on 10000 iterations.

## Supplementary data

**Fig. S1.**
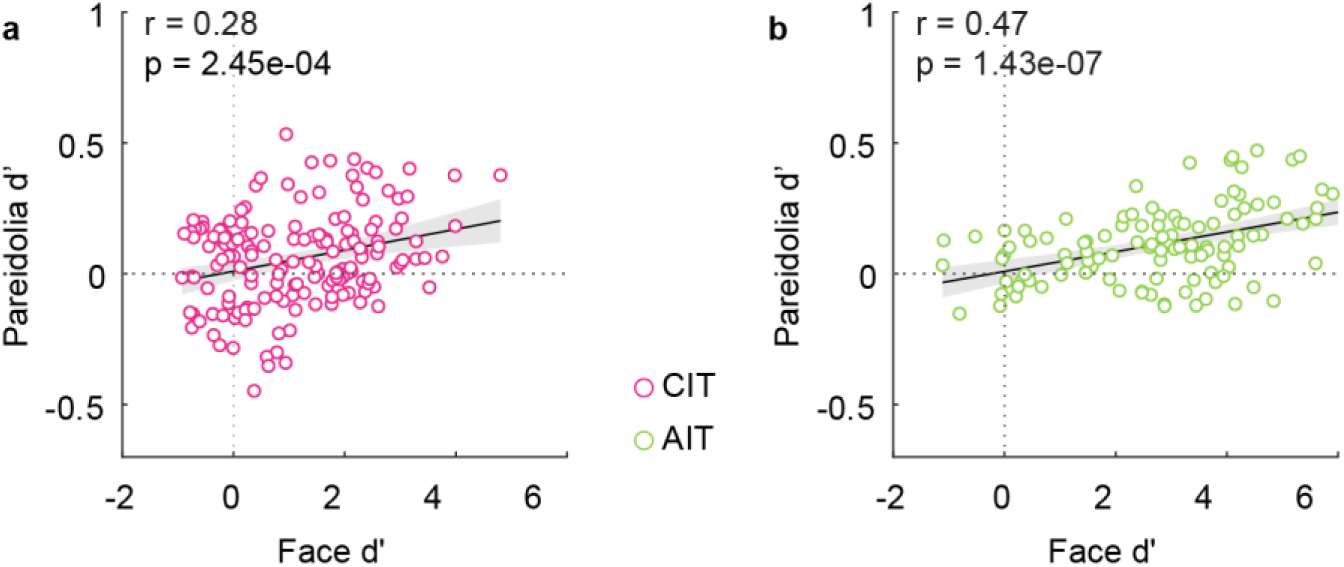
Pareidolia selectivity is not driven by image familiarity. **a.** Scatterplot showing the correlation between face selectivity (face d’) plotted on the x-axis and pareidolia selectivity (pareidolia d’) on the y-axis. Note that for this experiment, the monkeys only saw quadrant scrambled versions of unfamiliar pareidolia images, without seeing the original versions. Each dot depicts a neural unit in CIT (n = 162). **b.** Scatterplot showing the correlation between face and pareidolia selectivity in AIT (n = 114). Same conventions as in **a**.

**Fig. S2.**
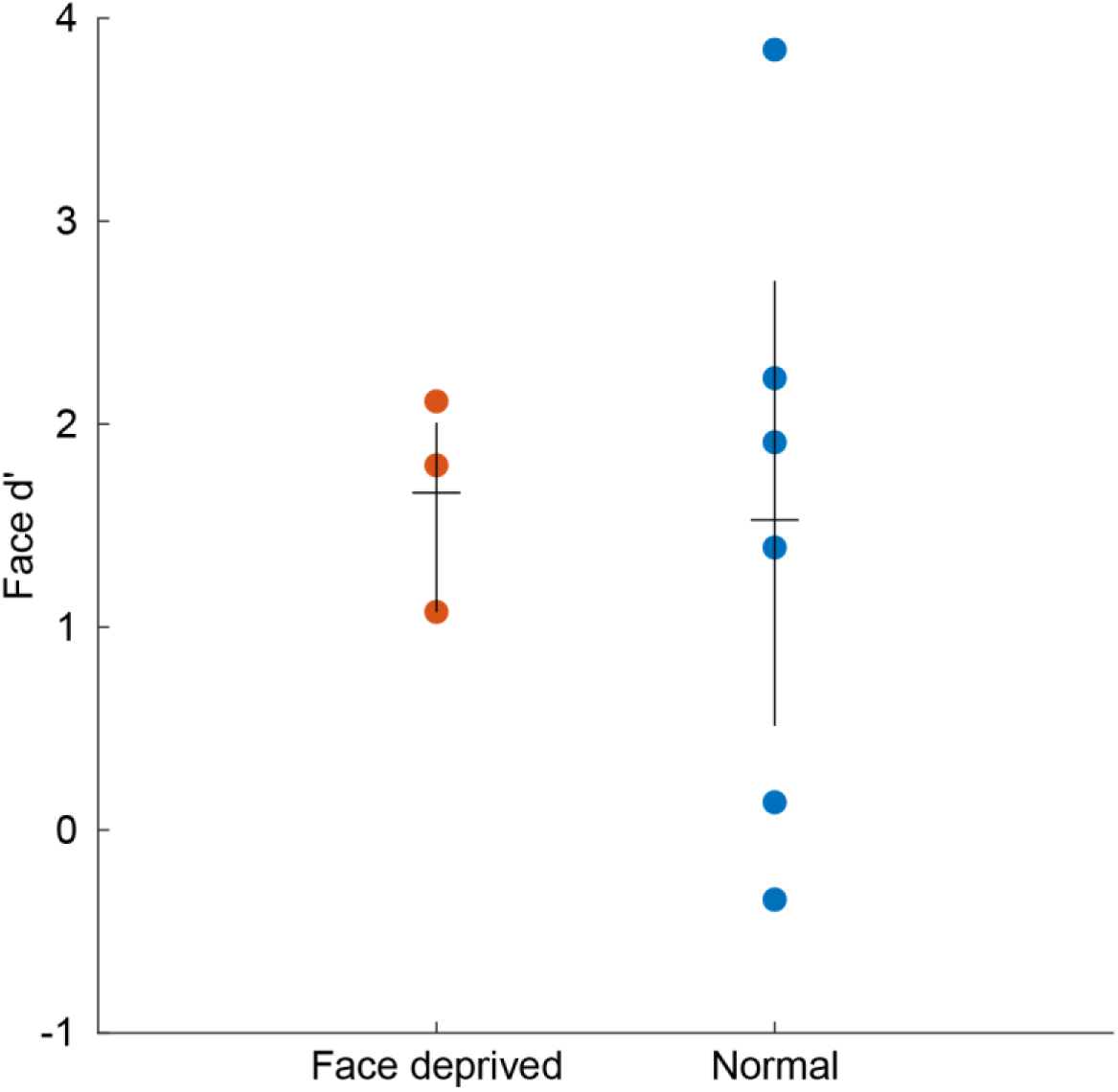
Face deprived and normal monkeys do not differ in face selectivity. Scatterplot showing the face selectivity (quantified by the metric face d’) on the x-axis for face-deprived (orange circles) and control monkeys (blue circles). The horizontal line shows the mean face d’ for each group, vertical lines depict the confidence intervals. The difference between face d’ for the face deprived monkeys was not significantly different from the control monkeys (t_7_ = 0.08, p = 0.94).

**Table S1.** Correlation between pareidolia and face selectivity per monkey. Table showing Pearson’s correlation *r* between pareidolia d’ and face d’ for each monkey in central (n = 4) and anterior IT (n = 5) for original and quadrant scrambled images from Experiment 1. FD = face deprived.

| Monkey ID (no. of units) | CIT |  |  |  | AIT |  |  |  |
| --- | --- | --- | --- | --- | --- | --- | --- | --- |
|  | Original |  | Scrambled |  | Original |  | Scrambled |  |
|  | r | p | r | p | r | p | r | p |
| M1 (64) | 0.53 | $8.4 \times 10^{-6}$ | 0.13 | 0.3 | | | | |
| M2 (FD; 73) | 0.63 | $1.9 \times 10^{-9}$ | 0.45 | $7.1 \times 10^{-5}$ | | | | |
| M3 (31) | 0.4 | 0.027 | 0.27 | 0.15 |  |  |  |  |
| M4 (FD; CIT: 40; AIT: 9) | 0.24 | 0.14 | 0.04 | 0.83 | 0.69 | 0.042 | 0.74 | 0.024 |
| M5 (60) |  |  |  |  | 0.44 | 0.0004 | 0.32 | 0.013 |
| M6 (13) |  |  |  |  | 0.74 | 0.0036 | 0.4 | 0.18 |
| M7 (24) |  |  |  |  | 0.58 | 0.0028 | 0.71 | 0.0001 |
| M8 (57) | | | | | 0.53 | $1.9 \times 10^{-5}$ | 0.4 | 0.0028 |
| Mean | 0.45 |  | 0.22 |  | 0.6 |  | 0.51 |  |

## Acknowledgments

We thank Dr Jessica Taubert and Dr Susan Wardle for sharing the stimuli. We also thank Dr Akshay Jagadeesh for helpful feedback and comments on the manuscript. This work was supported by the Harvard Lefler Fellowship, Hearst Fellowship and Gordon Fellowship (to S.S.), Alice and Joseph Brooks Fund Postdoctoral Fellow (K.V.), NIH grant EY16187 and P30 EY012196 and R01 EY025670 (to M.S.L.).

